# Pangaea: A modular and extensible collection of tools for mining context dependent gene relationships from the biomedical literature

**DOI:** 10.1101/2020.04.02.022517

**Authors:** Liviu Pirvan, Shamith A. Samarajiwa

## Abstract

**Motivation:** Pangaea is a scalable and extensible command line interface (CLI) software that integrates gene-relationship detection features to extract context-dependent structured gene-gene and gene-term relationships from the biomedical literature. It provides computational methods to identify biological relationships between a collection of genes and can be used to search and extract different types of contextual relationships amongst genes.

**Results:** We implemented a CLI-based software for downloading PubMed articles and extracting gene relationships from abstracts using natural language processing methods. In terms of scalability, the software was designed to support the retrieval and processing of millions of articles whilst minimising memory requirements and optimising for parallel processing on multiple CPU cores. To allow extensibility, the tool permits the use of contextual custom-made models for the text processing parts, and the output is serialised as JSON objects to allow flexible post-processing workflows.

**Availability:** The software is available online at: https://github.com/ss-lab-cancerunit/pangaea

## Introduction

Owing to the exponential growth of biomedical publications in databases such as PubMed (approximately 30 million articles in the first quarter of 2020), it is increasingly difficult for biomedical researchers to read or keep up with all the articles relevant to a particular field of study. Concurrently, the magnitude of literature constitutes a challenge in terms of computational resources required for text processing. As such, Natural Language Processing (NLP) based methods for extracting biological relationships between genes has been an area of active research[4], leading to many computational methods and webtools such as Chilibot [1], GeneCards [6] and the now defunct iHOP [2]. However, there is a distinct lack of high-throughput computational gene-relationship extraction methods with flexible and extensible functionality available, and Pangaea addresses this shortcoming.

Pangaea is a Python-based package with a CLI that provides an offline programmatic tool that enables text processing of scientific articles, applying text mining methods to structure relevant information. As such, the tool may be employed for use-cases such as retrieving context specific interactions among a known set of genes, identifying interactions between known genes and unknown targets, as well as confirming known relationships for pairs of genes. Along with the main software, we provide a set of tools for processing and exploring the extracted information.

## Approach

Pangaea is designed to prioritise extensibility and scalability of information extraction from biomedical literature. As such, its components are modularised to solve a set of loosely coupled tasks. For an overview of the system components and workflow, please refer to Figure 1.

**Fig. 1.**
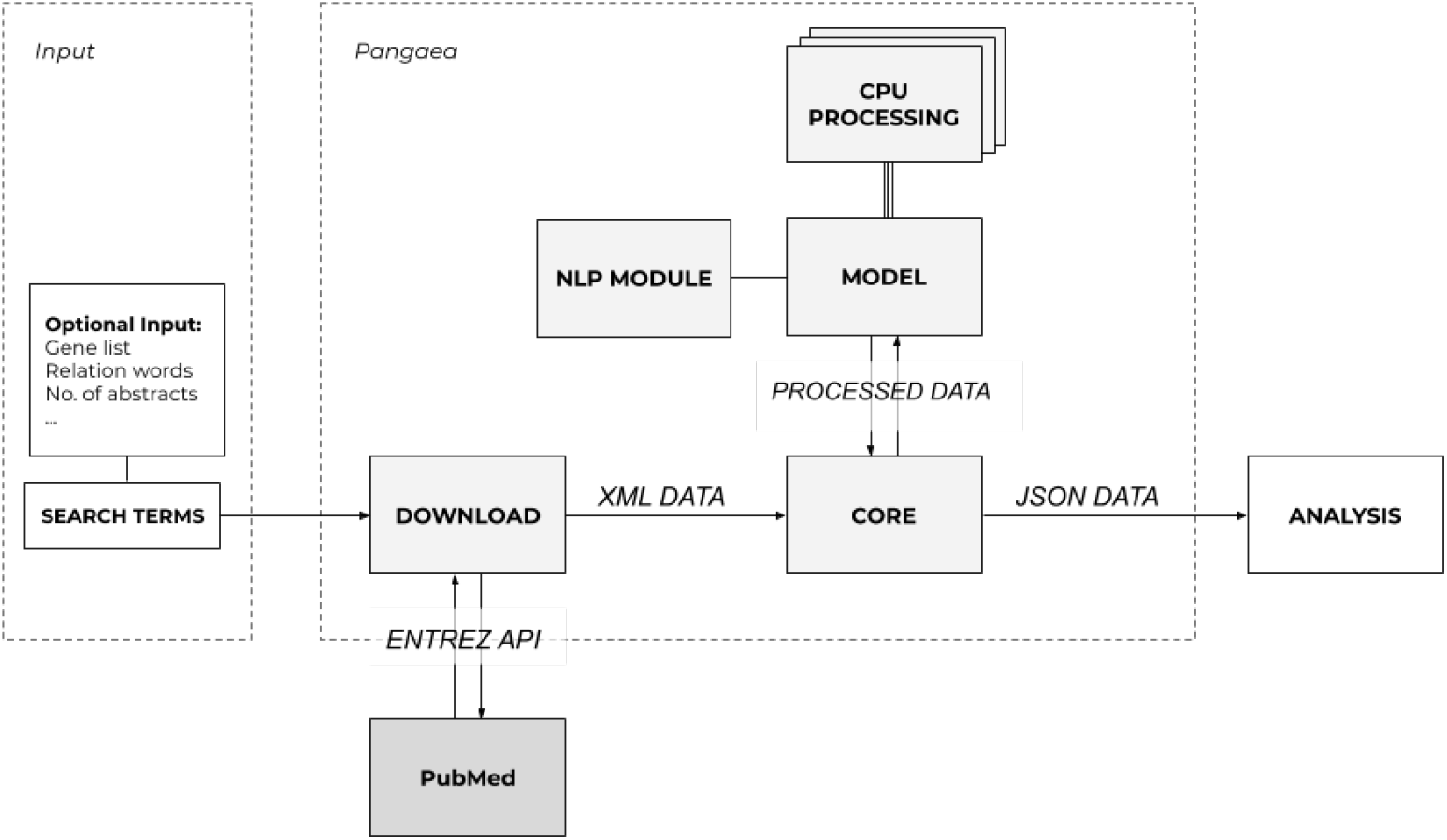
Diagram representing the standard workflow of using Pangaea.

The first component of the system is a tool wrapping the Entrez e-utilities [7] to communicate with the PubMed database and retrieve relevant abstracts. When provided with a search term and a value denoting the maximum number of abstracts to be retrieved, the data is downloaded locally and merged into an XML file that serves as input for the processing stage. According to the guidelines provided by Entrez and NCBI [7], we implement mandatory delays and batching of requests to lower the computational and bandwidth requirements. Lastly, the user must ensure that there is sufficient available storage to store the number of abstracts and relevant metadata required.

Then, the data downloaded serves as input for the processing stage. To avoid prohibitive memory requirements, the processing of the data is performed incrementally, discarding articles from memory once processed. Further, the processing of each abstract is independent from each other, so each article is submitted to a process pool shared among the CPUs. Once the processing of an article is completed, the output is stored in a process-shared queue. Writing to disk is accomplished asynchronously, writing the contents of the queue and flushing the buffer periodically to save progress and balance I/O demands. Thus, the system is designed to maintain constant memory requirements with respect to the size of the input file, and computationally it scales approximately linear with respect to the number of CPUs used.

To process the data, we use models based on natural language processing principles along with additional information regarding gene synonyms and keywords related to relationships between genes. For this step, the two main tasks are named-entity recognition (NER) for gene detection and entity-relationship extraction for identifying relationships between the genes. A customised relationship term list is supplied by the user, enabling customisation of the search domain. A suitable model may be highly dependent on the task that is required: for instance, the performance of the model can be improved using deep learning and word embeddings [3] at the cost of processing speeds and computational power required. Therefore, we designed our code base to provide an interface class that allows users to implement custom models if necessary. For the example results discussed in this paper, we used a model based on traditional NLP techniques including tokenization, lemmatization, parts-of-speech (POS) tagging, and n-grams [5].

## Methods

The software is developed as a CLI-based tool written in python3 for Linux Fedora systems, the code is hosted on GitHub, and it can be installed using the pip package-manager system.

The explicit inputs required are either search terms and a relationship dictionary to retrieve the data from PubMed or a file containing previously downloaded abstracts (XML) that can be processed. If the data is retrieved from PubMed, the tool adheres to the guidelines outlined for their API usage^1^, which recommends no more than three network requests per minute. Although downloads are batched according to the API guidelines, the retrieved files are merged locally into one XML file using sed before processing the contents using python3. With respect to performance, the processing time varies according to the task requirements and resources allocated for the tool (see Figure 2). Once the processing is complete, the output is a single JSON file containing a list of articles comprised of meta-information such as the title of the article and the journal in which it was published, as well as the gene relation sentences that were extracted.

**Fig. 2.**
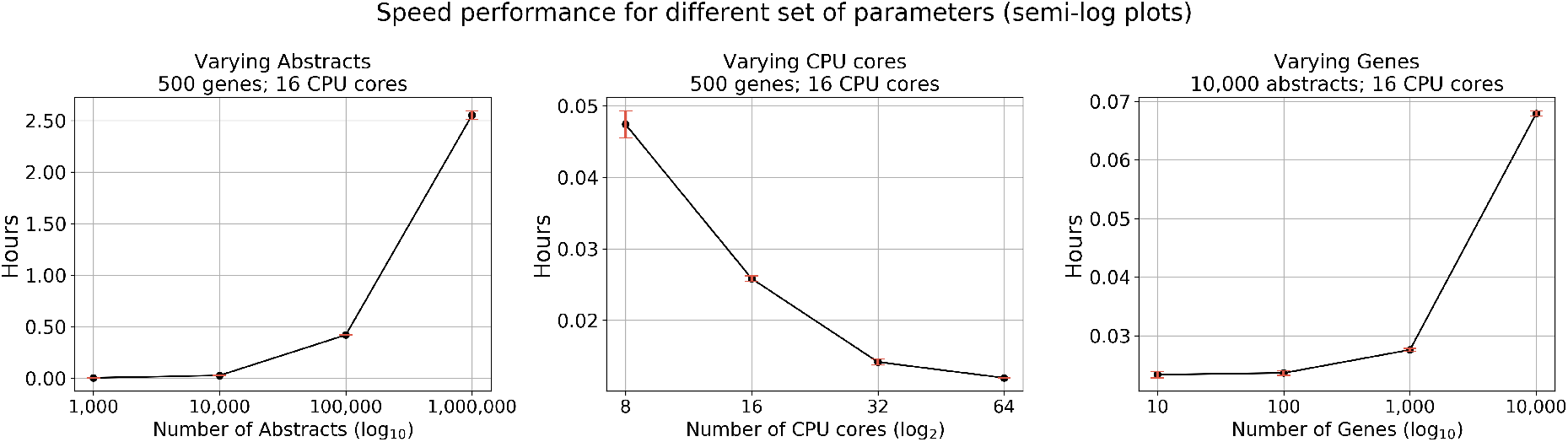
Performance graphs while varying number of genes considered, number of CPUs used, and the number of abstracts parsed.

To extract relevant information from the JSON file we provide example code in the form of Jupyter notebooks (see Figure A.1 and Figur A.2 for example usage), as well as instructions to integrate the data in a local search engine based on Elastic Search. We chose JSON as the preferred serialization format as libraries for processing JSON format are widely available for popular programming languages, and users can decide to process the information to fit their requirements. For more details, online documentation is available at https://pangaea-tool.readthedocs.io/en/latest/ and the code is hosted on GitHub at https://github.com/ss-lab-cancerunit/pangaea.

## Conclusion

Pangaea is a biomedical relationship extraction system that can select abstracts relevant to a particular biomedical domain and extract gene-gene or gene-term relations conforming to a contextual set of relationships. This scalable and extensible system provides a new architecture for mining, integrating and discovering information in biomedical texts.

## A Supplementary Data

### A.1 Figures

**Fig. A.1.**
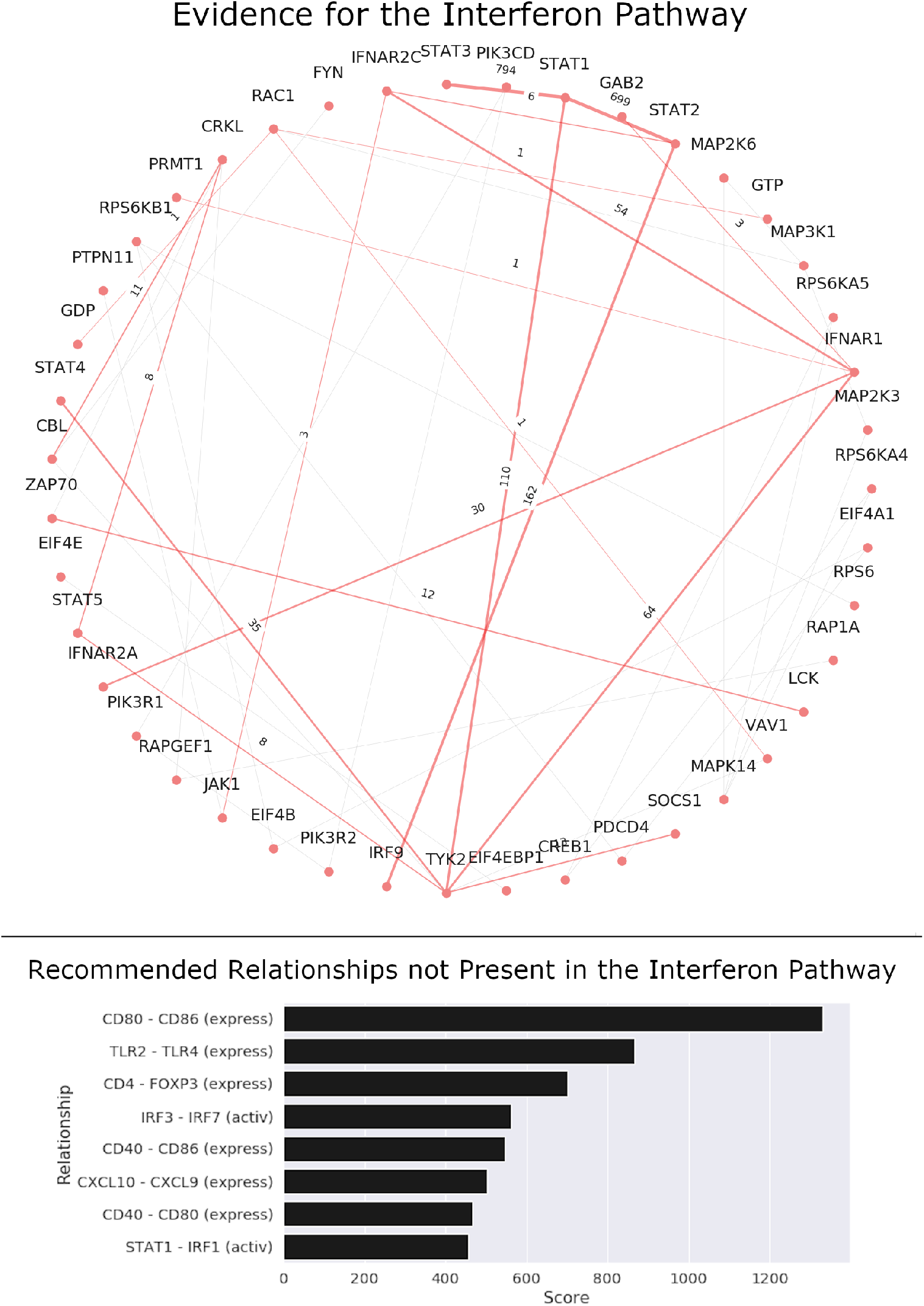
Pangaea output can be used with known pathway data to visualise evidence from the literature and provide recommendations for relationships absent from the pathway. In the top figure, a graph is used to visualise the degree of evidence for each interaction in the Interferon Pathway: the nodes are the genes analysed while the thickness of the edges encode the confidence in the interaction between the two genes. In the bottom figure, we display the top 8 relationships for which Pangaea found interaction evidence despite not being in the pathway provided.

**Fig. A.2.**
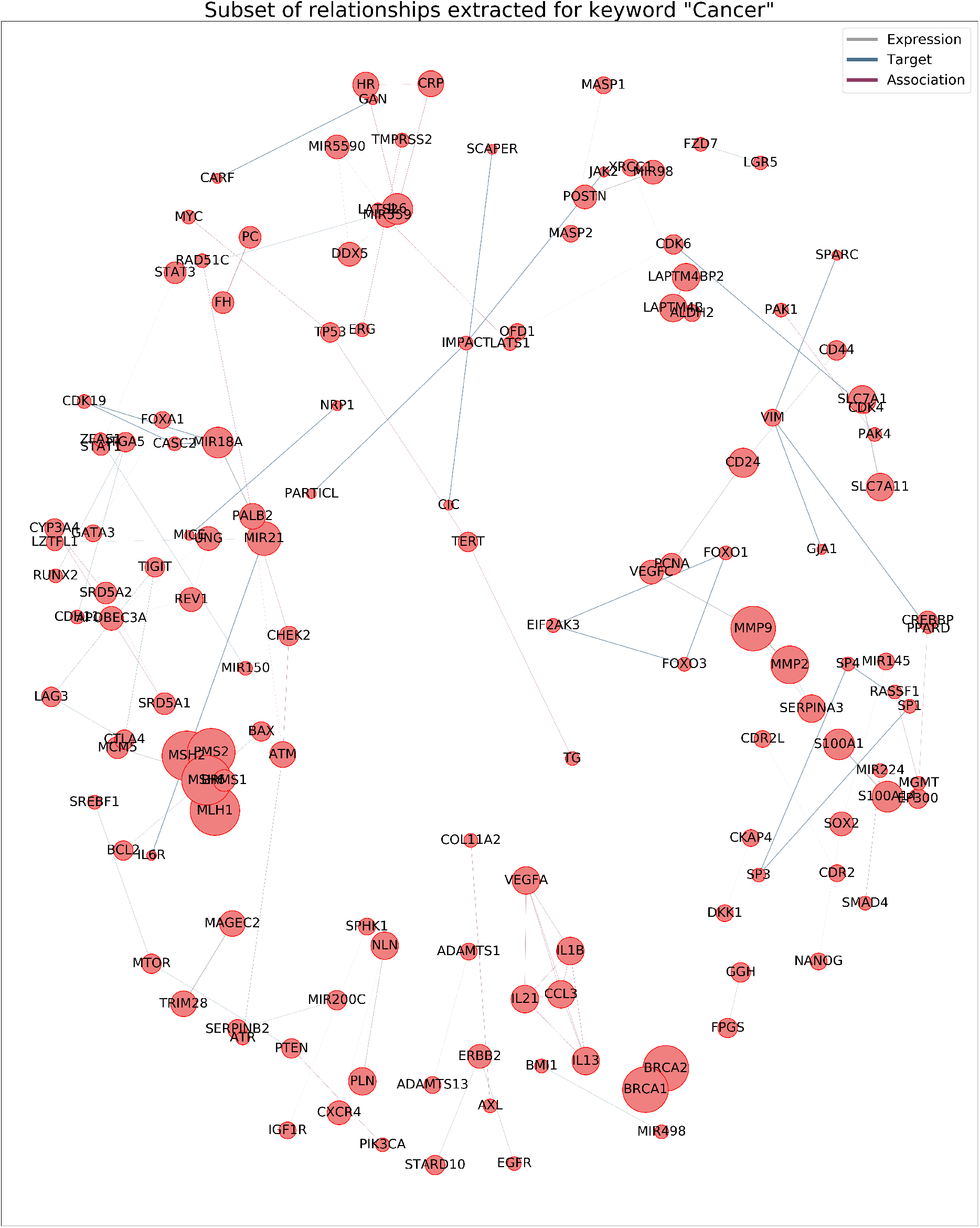
The graph above visualises a subset of gene interactions found by Pangaea for the keyword “cancer”.

### Footnotes

1 Please read this page for more information regarding usage guidelines: https://www.ncbi.nlm.nih.gov/books/NBK25497/

